# The variant catalogue pipeline: A workflow to generate a background variant library from Whole Genome Sequences

**DOI:** 10.1101/2022.10.03.508010

**Authors:** Solenne Correard, Mohammed OE Abdallah, Brittany Hewitson, Wyeth W. Wasserman

**Affiliations:** BC Children’s Hospital Research Institute, University of British Columbia, Vancouver BC, Canada; Centre for Molecular Medicine and Therapeutics, Department of Medical Genetics, University British Columbia, Vancouver BC, Canada

## Abstract

Today, several projects are working toward reducing inequities and improving health care for individuals affected with rare genetic diseases from diverse populations. One route to reduce inequities is to generate variant catalogues for diverse populations. To that end, we developed the variant catalogue pipeline, an open-source pipeline implemented in the Nextflow framework. The variant catalogue pipeline includes detection of single nucleotide variants, small insertions and deletions, mitochondrial variants, structural variants, mobile element insertions, and short tandem repeats. Sample and variant quality control, allele frequency calculation (for whole and sex-stratified cohorts) and annotation steps are also included, delivering vcf files with annotated variants and their frequency in the cohort. Successful application of the variant catalogue pipeline to 100 publicly available human genomes is described. We hope that, by making this pipeline available, more under-represented populations benefit from enhanced capacity to generate high-quality variant catalogues.

## INTRODUCTION

Next Generation Sequencing allows diagnostic success and health care management for patients affected with rare genetic diseases. However, not every patient benefits from these improvements equally. Indeed, most genomic research has been performed on populations of European ancestry and therefore benefits them more [1]. Several projects around the world are working toward reducing inequities and improving health care for individuals affected with rare genetic diseases from diverse populations [2,3].

One route to reduce inequities in genomic health care is to generate variant catalogues for diverse populations to more accurately identify and annotate variants implicated in rare genetic diseases. The main variant catalogue to date is gnomAD, with 76,156 genomes in the v3.1 short variant dataset [4]. While there is commitment to addressing diversity, some populations remain absent or barely represented within gnomAD. Hence, parallel genomic data generation projects have been pursued, focusing on specific populations, such as the Iranian population [5], the Korean population [6], the Polish population and many more.

Recently, several projects were launched with a special emphasis on Indigenous populations, such as the Silent Genomes Project (focusing on the Indigenous populations of Canada) and the Aotearoa Variome project (focusing on the Māori population in New-Zealand) [2]. Other projects, such as the All Of Us Research Program in the USA, also sought to include Indigenous populations. However, they have faced challenges related to the engagement with and governance of data pertaining to Indigenous peoples [8,9].

To enable such efforts, it is imperative to create and streamline pipelines for the processing of human genome sequence data. Enterprise-scale pipelines such as the one developed by the gnomAD team are extensive and publicly available [10]. However, the pipeline developed by the gnomAD team is highly complex and generally geared toward cloud platforms rather than local high-performance computing systems, thus limiting their portability for projects restricted by privacy laws or data sovereignty requirements. Moreover, addition of variant class detection methods over time (mitochondrial variants, STR, etc) has resulted in the creation of disparate pipelines. Thus, a need for a low barrier, shared, reusable comprehensive pipeline that incorporates leading-edge tools remained unfulfilled.

To that end, we developed the variant catalogue pipeline, an open-source pipeline implemented in the Nextflow framework [11], that can be used on local servers, and is modular to allow each project to use as much or as little of the pipeline as they wish. We hope that, by making this pipeline available, more under-represented populations around the world will have capacity to generate high-quality variant catalogues. The variant catalogue pipeline includes detection of Single Nucleotide Variants (SNV), small insertions and deletions (indels), Mitochondrial variants, Structural Variants (SV), Mobile Element Insertions (MEI), and Short Tandem Repeats (STR). The variant catalogues can be generated for GRCh37 and/or GRCh38 human reference genomes.

## MATERIAL AND METHODS

The variant catalogue pipeline is an automated workflow that takes as input Whole Genome Sequence (WGS) short-reads data (fastq) and outputs multiple vcf files including the variant allele frequencies in the cohort and some basic annotation (Overview in Figure 1, detailed in supplementary figure 1).

**Figure 1.**
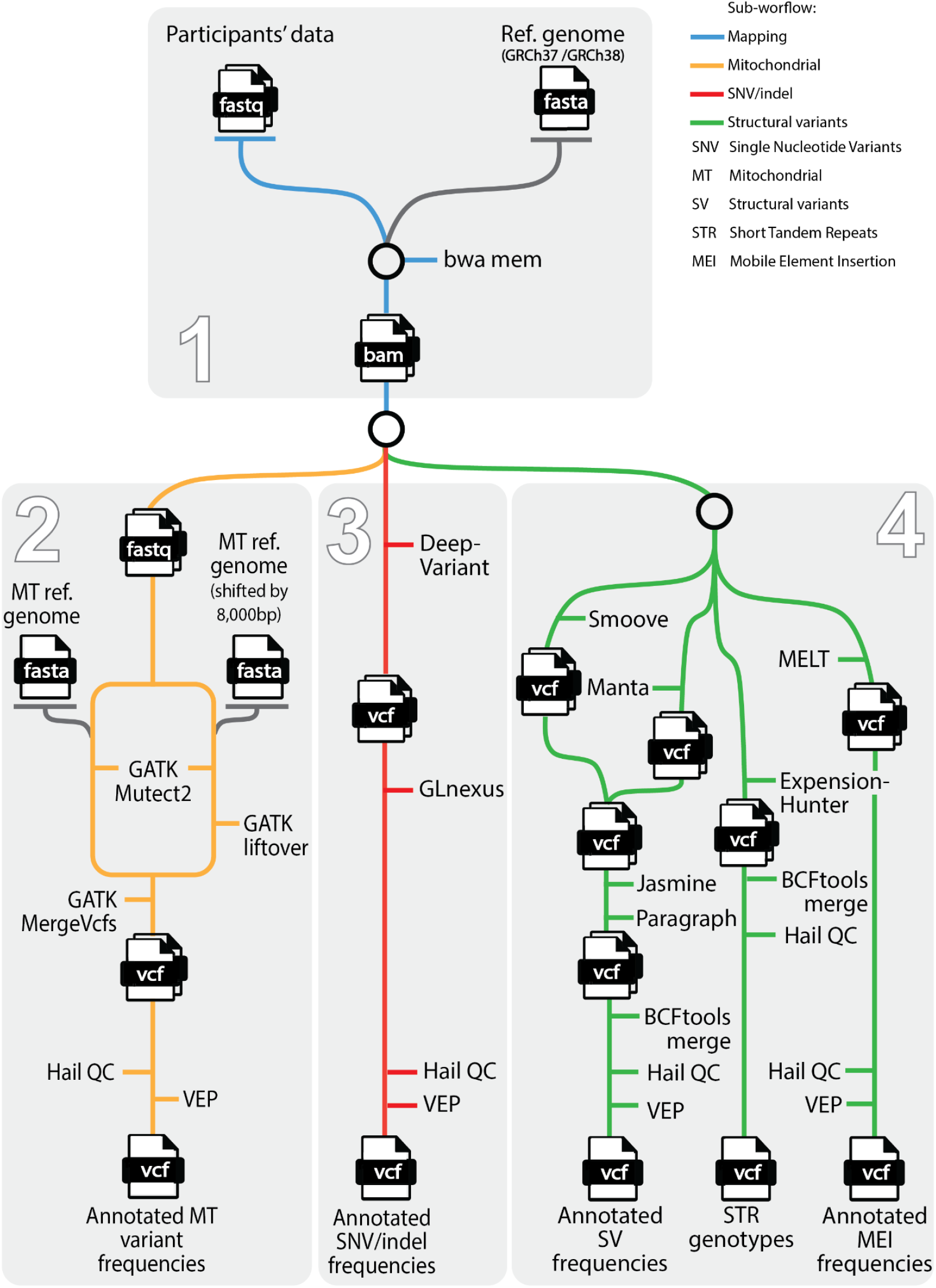
Overview of the variant catalogue pipeline. Each color represents a sub-workflow (blue: mapping, yellow: mitochondrial, red: SNV/indels, green: structural variants). A more detailed version of the variant catalogue pipeline is available as a supplementary figure.

The variant catalogue pipeline is implemented in the Nextflow framework to allow modularity for future users, portability of the pipeline to different systems, reproducibility, and tracking of error logs and intermediate files. The variant catalogue workflow consists of four sub-workflows :

1. Mapping sub-workflow (detailed in supplementary data) takes as input a reference genome and two fastq files per individual, and outputs one bam file per individual, as well as a quality overview of the individual data (based on quality control of the fastq and bam files, html file).
2. SNV and indel sub-workflow (detailed in supplementary data) takes as input a reference genome and bam files (one per individual) and outputs one vcf file with filtered annotated variant frequencies for the SNV and indels identified in the cohort. Several intermediate files are generated, including individual vcf files, cohort vcf files with the genotype of each individual for each variant, before and after variant filtration.
3. Mitochondrial variant sub-workflow (detailed in supplementary data) takes as input two mitochondrial reference genomes (GRCh38 MT reference genome and the same reference genome shifted by 8,000bp) and bam files (one per individual) and outputs a vcf file with filtered annotated mitochondrial variant frequencies identified in the cohort. Several intermediate files are generated, including individual vcf files, cohort vcf files with the genotype of each individual for each variant, before and after variant filtration.
4. Structural variant sub-workflow (detailed in supplementary data) takes as input bam files (one per individual) and outputs vcf files with filtered annotated structural variant frequencies identified in the cohort. Several intermediate files are generated, including individual vcf files, and cohort vcf files including the genotypes of all sequenced individuals before and after variant filtration.

Each sub-workflow is independent and can be run individually if the requisite input files are available. For example, it is possible to run only the SNV and indel sub-workflow if the bam files for each individual of the cohort are available, even if the bam files were generated using an alternative method from the one in the mapping sub-workflow. Moreover, each sub-workflow outputs useful files at several stages along the process, hence, it is not necessary to use a complete workflow if a user seeks an intermediate file, such as the unfiltered cohort vcf file. Each sub-workflow is composed of modules, which call upon open-access genomic software.

## RESULTS

In order to test the variant catalogue pipeline, we processed 100 samples from the IGSR (International Genome Sample Resource, [12]). The input data is paired-end whole genome sequences (fastq) with an expected average coverage of 30X and the output data are the vcf files containing the variant frequencies and annotations. The input files are publicly available (see data availability section).

To simulate samples coming in several batches, Batch 1 was composed of 80 samples, and Batch 2 was composed of 20 samples. The steps that are “individual specific” (above the dashed line in supplementary figure 1) don’t need to be reprocessed if the sample was part of a previously processed batch.

The pipeline was processed on a local high-performance computing system using three compute nodes (40 CPUs/node and 384G RAM/node) managed by SLURM scheduler. From start to finish, the processing of 80 samples (batch 1) took about 26,000 CPU-hours. Processing of the second batch containing 20 additional samples to obtain the variant frequencies among the cohort of 100 individuals took 14,300 CPU-hours, the detailed report generated by Nextflow is available in the GitHub repository (html format).

For the 100 samples, the initial data size is 5.7 Tb, the size of the final vcf files combined is less than 35 Gb while the total size of the output, including the bam files, several vcf files, and other intermediate files is less than 5.5 Tb.

Finally, post-processing quality control steps performed in Hail identified nine outlier samples (9%) [13]. The filtered output vcf files included a total of 23,976,026 SNV/indel, 1,567 Mitochondrial variants, 117,071 structural variants and 6,807 MEI.

## DISCUSSION

While variant catalogues remain a major goal for many communities, obstacles hinder this pursuit. One of the major challenges is the design and choice of pipelines and workflows to process the genomic data from whole-genome sequence data to annotated DNA sequence variants and their shareable allele frequencies. The variant catalogue pipeline offers a comprehensive pipeline to identify variants and calculate allele frequencies within a cohort of whole genome sequences.

### Comparison of the existing variant catalogue pipelines

One of the most extensive and widely used variant catalogue to date is gnomAD [4]. Their variant processing pipeline has been configured to be executed on Google Cloud via the Workflow Description Language (WDL) and the Cromwell Execution Engine and remains unusable for most projects in lieu of its esoteric complexity and its reliance on commercial cloud platforms. As we couldn’t process the same subset of data with the gnomAD pipeline, we offer a broad comparison of the techniques and tools used in both pipelines.

The mapping steps of the variant catalogue pipeline are highly similar to that of gnomAD. The mitochondrial sub-workflow, as well as the mitochondrial variant quality control steps, reuse parts of the gnomAD code [14]. The STR genotyping part is highly congruent with gnomAD, both using ExpansionHunter and the 59 STR loci with disease associations [15].

The SNV sub-workflow differs from gnomAD as the variant catalogue pipeline relies on DeepVariant [16] (for SNV calling) and GLnexus [17] (for joint calling), while gnomAD relies on GATK HaplotypeCaller (for SNV calling) and Hail (for joint calling).

The structural variant sub-workflow also differs in various ways from gnomAD SV discovery methods [18]. gnomAD SV pipeline executes four independent SV detection algorithms per sample: Manta [19], DELLY [20], MELT [21], and cn.MOPS [22]. The variant catalogue pipeline executes three : Manta and MELT, which are common with gnomAD, and Smoove [23], which is unique to the variant catalogue pipeline. The following modules from the gnomAD pipeline (Variant filtering, Genotyping, Batch integration and vcf refinement) were not applicable as different variant callers were used. The details of the structural variants sub-workflow are available in supplementary data.

Another major difference between the variant catalogue pipeline and the gnomAD pipeline is the absence of ancestry inference, mitochondrial haplogroup assignment and relatedness calculation. While we acknowledge the relevance of such metrics for projects including large cohorts, we focused solely on allele frequency calculation and variant annotation. For Indigenous genetics, misuse of data in past projects has highlighted the central importance of Indigenous governance and control of Indigenous data [24]. We understand the scientific relevance of ancestry and relationship information to population-specific variation projects, such as understanding the frequency of alleles in sub-populations or the detection of samples with outlying characteristics to the population. Such analysis sub-workflows could be developed by projects seeking to include such information in their variation catalogue. Alternatively, it is also possible to do similar analysis in a manner that is respectful of project governance choices, such as the development of the allele dispersion score, which allows distinguishing between variants spread across a cohort and variants restricted to subsets of genetically more similar individuals [25]. This does not require classifying individuals into groups, nor does it require sharing population specific information.

Many variant catalogues generated to date, such as the H3Africa genome project [26], the Saudi Human Genome Program [27], the Qatari Genome project [28] or SweGen [29], use in-house custom scripts relying on GATK best practices for SNV and indel variant calling. More recent projects, like the Thousand Polish Genomes [30] used DeepVariant and GLnexus for SNV and indel calling, like in the variant catalogue pipeline. SweGen also included a structural variant analysis relying on Manta 1.0.0.

### Comparison of the variant catalogue pipeline to other existing variant calling pipelines

Other projects sought to develop open-source pipelines for project processing whole genomes data. A particularly close project is the Sarek pipeline designed to detect variants on whole genome or targeted sequencing data [31]. While many modules overlap with the variant catalogue pipeline, it lacks the comprehensive coverage of structural variant classes and the key focus on background variant library generation.

The Swedish Genome project is developing a Nextflow pipeline for rare disease diagnosis using WGS and RNA-seq data, however, it remains unpublished and lacks features provided in the variant catalogue pipeline. Similar pipelines are available from commercial vendors such as Illumina Dragen and Sentieon DNASeq, but the costs can be prohibitively costly for use at population-level projects.

### Data sovereignty : The necessity to be able to bring tools to data

The Silent Genomes project, for which the work described in this manuscript was generated, is developed in partnership with Indigenous leadership, communities and individuals. Indigenous leadership and governance over the production and access of Indigenous genomic data is central to the project [32]. Due to concerns related to data sovereignty, it was decided to exclude cloud services and other large-scale computing servers. Processing large amounts of genomic data on local servers can be a challenge requiring large storage and computing capacity, as well as system management to install specialized software, hence bringing tools to data can be challenging for processing of Whole Genome Sequences on larger scales. While the workflow provided here may help, future development in genomic research will need to continue progress towards enabling options for local control of sensitive data.

### Future extension of the variant catalogue pipeline

The sequencing technologies keep evolving over time, and we hope that the pipeline will evolve in parallel. For example, the increasing interest in long-read sequencing technology will require the development of a mapping sub-workflow specific to long-read data. Moreover, we appreciate and acknowledge that many genome analysis tools exist and many overlap in purpose (i.e. SNV calling). Our pipeline is modular and a user can readily modify modules to use the alternative software of their choosing. Finally, researchers are developing variant libraries for diverse mammalian species [33, 34]. Even though this pipeline was developed for the human species, it would be easily adaptable to other mammalian species for which there is an available reference genome.

To date, two techniques are used by the scientific community to share variant catalogues: through an interactive interface and / or downloadable vcf files. The variant catalogue pipeline generates vcf files containing the allele frequencies across the cohort that, if the project allows, can be deposited online for the scientific community to download and use.

Working towards greater equity in genomics, we developed a complete genetic variation data pipeline, the variant catalogue, suitable for both cloud and local computation, to enable population-specific projects to generate data across variant classes. Most population-specific catalogues only report the frequencies of SNVs and indels, and this pipeline will allow them to expand the variant classes they report. In sharing the pipeline, we hope to encourage long-term cooperation between population-specific projects requiring low-barrier pipeline access and for the community to build on it and maintain it so that future projects can deliver reliable variant catalogues for every population.

## AVAILABILITY

### Pipeline

The variant catalogue pipeline is implemented in the Nextflow framework and relies only on open-access tools, therefore, any user with sufficient compute capacity should be able to use this pipeline. Users who want to use this pipeline on their local servers will have to install the necessary software on their instance.

The variant catalogue pipeline is available in the Wasserman lab GitHub repository https://github.com/wassermanlab/Variant_catalogue_pipeline. For clarity, it is organized in sub-workflows and modules as presented in this article.

All the softwares and other resources necessary to run the variant catalogue pipeline (Reference genomes, annotation plugins, etc) are open-source and the information related to them is available in Github under : https://github.com/wassermanlab/Variant_catalogue_pipeline/tree/main/supplementary_information.

### Test case : 100 genomes from the IGSR

The details to download the 100 genomes used in this test, as well as the script to pre-process them (from cram to fastq), are available on GitHub under : https://github.com/wassermanlab/Variant_catalogue_pipeline/tree/main/test_case.

Output files generated by Nextflow (report, timeline, etc) together with vcf files containing annotated variants and their frequencies by variant type are also available in the same repository. Intermediate files were not uploaded to GitHub because of their size.

## ACKNOWLEDGEMENT

The authors acknowledge the lands on which they work, live and play: the traditional, ancestral, and unceded territory of the xʷməθkʷəýəm (Musqueam), Skwx̱ wú7mesh (Squamish), and səlilwətaɬ (Tsleil-Waututh) Nations. We gratefully acknowledge past and present members of the Wasserman lab and the Silent Genomes Project team for helpful discussion. We thank Irina Manokhina and Dora Pak for project management support.

## Supplementary Data

### Details of the mapping sub-workflow

In this initial workflow step, a reference genome and two fastq files per individual are input, and one bam file per individual are output. The sub-workflow works with paired-end short-read sequences but could be adapted to support long-read sequences using appropriate mapping software. This sub-workflow can be skipped if bam files for individual genomes are available.

The short-read mapping is performed using bwa mem with standard parameters [1], the generated sam file is compressed, sorted and indexed using samtools [2] to obtain an indexed bam file per individual.

This sub-workflow also includes quality control analysis of the individual data (fastq and bam). FastQC is used with the fastq files as input [3]. Mosdepth is used with the bam files as input to define the average coverage of each sample and the coverage of each chromosome [4]. Three Picard options are used (CollectWGSMetrics, CollectAlignmentSummaryMetrics and QualityScoreDistribution) with bam as input [5]. The output of the five software tools are aggregated using multiQC [6], outputting an interactive html file, allowing users to identify outliers.

### Details of the SNV and indel sub-workflow

The Single Nucleotide Variant (SNV) and indel (short insertions / deletions) sub-workflow takes as input individual indexed bam and outputs vcf files with filtered annotated variant frequencies for the SNV and indels identified in the cohort.

The variant caller used is DeepVariant, using reference parameters [7]. It outputs individual vcf files and gvcf files. Then GLnexus, with default parameters, is used to perform the joint calling of the individual vcf file and output one bcf with aggregated information for all samples [8]. Finally, bcftools is used to split the multiallelic variants (norm -m), rename the variants (annotate --set-id) and compress the vcf file [2]. GATK IndexFeatureFile is used to index the compressed vcf file [9].

This vcf file is then loaded into Hail [10], a tool developed to process quickly large amounts of genomic data, to filter samples and variants based on quality data. The first Hail step (sample QC) identify outliers and defines the sex of each sample (XX, XY or ambiguous) based on the genomic information only.

The sample quality control metrics were generated using Hail which computes summary statistics per sample. Samples falling quality control were determined as samples with any of the following QC metric falling outside cut-offs (cut-offs were set as three standard deviations of the mean, but can easily be changed to users preference) : approximate Read Depth (DP), Genotype Quality (GQ), fraction of calls neither missing nor filtered (call_rate), number of SNP alternate alleles (n_snp), number of private alleles (n_singleton), Insertion / Deletion allele ratio (r_insertion_deletion) and Transition / Transversion ratio (r_ti_tv).

The sex inference is identical to the one used by gnomAD for the genomes: the inbreeding coefficient (F) calculated through Hail is used, samples with F > 0.8 were classified as male, samples with F < 0.5 were classified as female and the other samples were classified as ambiguous.

The output is a list of samples that passed the quality control step and their genetic sex, as well as graphs and summary files with quality information for each sample.

The second Hail step (variant QC) filters out low quality variants as well as indels longer than 50bp. The variant quality control metrics were generated using Hail which computes variant statistics from the genotype data. Variants falling quality control were determined as variants with any of the following QC metric falling under the cut-off values (the cut-off values were set as three standard deviations under the mean, but can easily be changed to users preference) : approximate Read Depth (DP), Genotype Quality (GQ), fraction of calls neither missing nor filtered (call_rate), total number of called alleles (AN), and number of samples with a missing GT (n_not_called). It outputs a filtered vcf file, including individual genotypes, as well as a “frequency file” which includes only variant frequencies for variants that passed the quality control, within the cohort of samples that passed the sample QC. The generated vcf file is then annotated using VEP with the following options : --symbol, --biotype, --transcript_version, --tsl, --hgvs, --variant_class, --distance 0 (to avoid upstream_gene_variant or downstream_gene_variant consequences), --merged (use Ensembl and RefSeq cache), --use_transcript_ref (especially relevant when using the RefSeq cache), --check_existing, --var_synonyms and the following prediction tools : --polyphen b, --sift b, --plugin CADD, --plugin SpliceAI [11], which outputs annotated vcf files (one per chromosome).

### Details of the mitochondrial variant sub-workflow

The mitochondrial variant sub-workflow is developed specifically to call variants located on the mitochondrial DNA. The complexity of mitochondrial genome analysis necessitated the development of a specialized workflow. The workflow is modelled after the gnomAD mitochondrial variant calling pipeline, available in wdl [12]. Laricchia et al., highlighted the complexity of calling variants on the mitochondrial genome and the solutions developed to address that, including aligning the reads against a “shifted” reference mitochondrial genome, explaining the parallel steps in the mitochondrial sub-workflow, and the use of Mutect2 as the variant caller. This workflow takes as input the individual bam files, as well as two mitochondrial reference genomes and outputs a vcf file with the frequencies of the filtered annotated mitochondrial variants identified in the cohort.

For each sample, reads mapping to the mitochondrial genomes are extracted from individual bams (generated by the mapping workflow or input by the user) using GATK PrintReads (with --read-filter MateOnSameContigOrNoMappedMateReadFilter and --read-filter MateUnmappedAndUnmappedReadFilter option as per the gnomAD pipeline). Then, the bam file containing only the reads mapping to the mitochondrial genome is converted to a fastq file using GATK SamToFastq. Each fastq file is mapped against the mitochondrial reference genome and the shifted mitochondrial genome using bwa mem, duplicated reads are tagged using GATK MarkDuplicates, and newly obtained bams indexed using samtools. For each newly generated bam (two per individual), variants are called using GATK Mutect2 using the --mitochondria-mode and recommended parameters for mitochondrial variant calling, outputting a compressed vcf file. Variants called using the shifted reference genomes are off by 8,000 base pairs compared to the variants called using the mitochondrial reference genome, therefore, variant coordinates are shifted using GATK LiftoverVcf and the chain file associated to the shifted reference genome. Then, for each individual, the two vcf files are merged using GATK MergeVcfs to obtain one vcf file per individual with optimal variant calls for the entire mitochondrial genome, then bcftools is used to remove duplicated calls (bcftools norm --rm-dup) and index the vcf file. Multiallelic variants are split using GATK LeftAlignAndTrimVariants with the --split-multi-allelics option, remaining variants are filtered using GATK FilterMutectCalls with the --mitochondria-mode and GATK VariantFiltration to remove the variants known to be artifacts from the list provided by the GATK team.

Then, individual vcf files are loaded into Hail which merges the individual vcf files into one matrix table, and filters samples and variants based on quality control metrics. The filtration steps are modelled after the gnomAD Hail pipeline for MT variants, the main difference is the absence of haplogroup and population use in the variant catalogue pipeline. Samples failing quality control for the mitochondrial variant sub-workflow may differ from those failing quality control for the SNV/indel sub-workflow. The Hail module outputs a filtered vcf file which is then annotated using vep (with the same options as for the SNV/indel annotation presented in supplementary data), which outputs an annotated vcf file.

### Details of the structural variant sub-workflow

The Structural Variants (SV) sub-workflow is designed for more complex variants such as Structural Variants (SV) (including deletions, insertions, duplications, inversions and translocations), Short Tandem Repeats (STR) and Mobile Element Insertions (MEI). It takes as input the individual bam files, as well as specific annotation files depending on the type of variants in consideration and outputs several vcf files with filtered annotated structural variant frequencies identified in the cohort. Several intermediate files are generated, including individual vcf file, cohort vcf file with the genotype of each individual for each variant before and after variant filtration.

For the structural variants, two callers are used: Smoove and Manta. For Smoove, the structural variants are called using default parameters and the output bcf is compressed, sorted, and indexed using bcftools. For Manta, the process is similar with variants called with default parameters and the output vcf file is indexed using bcftools. Then, for each individual, the two vcf files are merged using bcftools concat to obtain one vcf file per sample. Then, the several vcf files are merged using Jasmine [13]. Each variant listed in the vcf file outputted by Jasmine is genotyped in each individual bam using Paragraph, outputting again one vcf file per individual [14]. Finally, these vcf files are merged using bcftools merge, and indexed using GATK IndexFeatureFile.

For the Mobile Element Insertions (MEI), MELT is used to identify mobile element insertions in each individual [15]. Then, the individual vcf files are merged using bcftools merge, and indexed using GATK IndexFeatureFile.

For the Short Tandem Repeats (STR), ExpansionHunter is used together with a pre-established STR catalogue containing 59 STR loci with disease associations to genotype the individuals. Then, the individual vcf files are merged using bcftools merge, and indexed using GATK IndexFeatureFile.

The three vcf files containing SV, MEI and STR are processed with an in-house Hail script for variant quality control: the structural variants smaller than 50bp are removed (to avoid overlap with short insertions/deletions called by the SNV pipeline), variants with a low quality tag in the vcf FILTERS field (such as MaxDepth, MinGQ, MinQUAL, etc), SV with an allele number (AN) value below the threshold of 75% of the maximum allele number (i.e. remove the variants that were not genotyped correctly in the cohort and will not give a reliable allele frequency estimation, the minimum threshold can easily be changed to users preference). Hail module outputs a file with the number of SV failing each filter. Given that there are no gold-standard benchmarking procedures for SVs from WGS, we validated the variant catalogue SV sub-workflow by visualizing a subset of SV using IGV [16].

For SV and MEI, samples falling sample quality control based on SNV genotype data (as described in supplementary data), were removed from the SV dataset before calculating the allele frequencies. The Hail module outputs for SV called by Smoove and Manta) and MEI are then annotated using vep (with the same options as for the SNV/indel annotation presented in supplementary data), which outputs an annotated vcf file per chromosome.

The STR output differs from the other variants, as the genotype of interest is the length of the repeat (allele size) and the genotype distribution. Therefore, for STR, the final output is an aggregated vcf file containing the individuals genotypes. Moreover, the annotation step is not done using VEP as the information relevant to each STR is included in a JSON file associated to the catalogue.

---

**Supp Figure 1.**
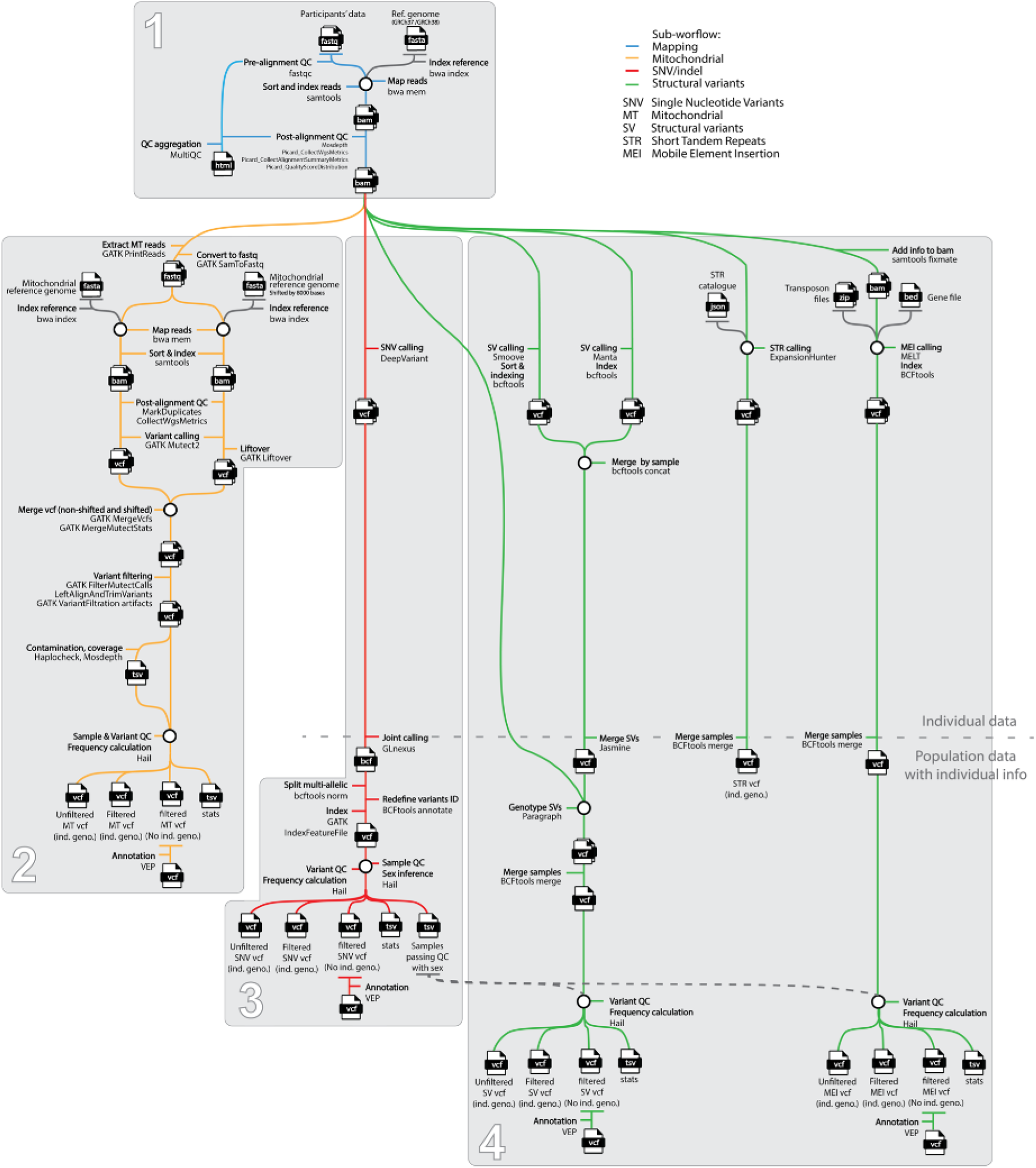
Details of the variant catalogue pipeline. Each color represents a sub-workflow (blue: mapping, yellow: mitochondrial, red: SNV/indels, green: structural variants). Each horizontal line represents a module within each sub-workflow. All the modules that are above the dashed line need to be run only once per sample, i.e. If a new batch of sample is included in the variant catalogue pipeline, it will not be necessary to reprocess the previously processed samples.

## REFERENCES

1. Landry, L.G., Ali, N., Williams, D.R., Rehm, H.L., and Bonham, V.L. (2018). Lack of diversity in genomic databases is A barrier to translating precision medicine research into practice. Health Aff (Millwood) 37, 780–785.

2. Caron, N.R., Chongo, M., Hudson, M., Arbour, L., Wasserman, W.W., Robertson, S., Correard, S., and Wilcox, P. (2020). Indigenous genomic databases: pragmatic considerations and cultural contexts. Front. Public Health 8, 111.

3. Appelbaum, P.S., Burke, W., Parens, E., Zeevi, D.A., Arbour, L., Garrison, N.A., Bonham, V.L., and Chung, W.K. (2022). Is there a way to reduce the inequity in variant interpretation on the basis of ancestry? Am. J. Hum. Genet. 109, 981–988.

4. Karczewski, K.J., Francioli, L.C., Tiao, G., Cummings, B.B., Alföldi, J., Wang, Q., Collins, R.L., Laricchia, K.M., Ganna, A., Birnbaum, D.P., et al. (2020). The mutational constraint spectrum quantified from variation in 141,456 humans. Nature 581, 434–443.

5. Fattahi, Z., Beheshtian, M., Mohseni, M., Poustchi, H., Sellars, E., Nezhadi, S.H., Amini, A., Arzhangi, S., Jalalvand, K., Jamali, P., et al. (2019). Iranome: A catalog of genomic variations in the Iranian population. Hum. Mutat. 40, 1968–1984.

6. Lee, S., Seo, J., Park, J., Nam, J.-Y., Choi, A., Ignatius, J.S., Bjornson, R.D., Chae, J.-H., Jang, I.-J., Lee, S., et al. (2017). Korean Variant Archive (KOVA): a reference database of genetic variations in the Korean population. Sci. Rep. 7, 4287.

7. Kaja, E., Lejman, A., Sielski, D., Sypniewski, M., Gambin, T., Dawidziuk, M., Suchocki, T., Golik, P., Wojtaszewska, M., Mroczek, M., et al. (2022). The Thousand Polish Genomes-A Database of Polish Variant Allele Frequencies. Int. J. Mol. Sci. 23.

8. Fox, K. (2020). The Illusion of Inclusion - The “All of Us” Research Program and Indigenous Peoples’ DNA. N. Engl. J. Med. 383, 411–413.

9. All of Us Research Program Investigators, Denny, J.C., Rutter, J.L., Goldstein, D.B., Philippakis, A., Smoller, J.W., Jenkins, G., and Dishman, E. (2019). The “All of Us” Research Program. N. Engl. J. Med. 381, 668–676.

10. Karczewski, K.J., Francioli, L.C., Tiao, G., Cummings, B.B., Alföldi, J., Wang, Q., Collins, R.L., Laricchia, K.M., Ganna, A., Birnbaum, D.P., et al. (2019). Variation across 141,456 human exomes and genomes reveals the spectrum of loss-of-function intolerance across human protein-coding genes. BioRxiv.

11. Di Tommaso, P., Chatzou, M., Floden, E.W., Barja, P.P., Palumbo, E., and Notredame, C. (2017). Nextflow enables reproducible computational workflows. Nat. Biotechnol. 35, 316–319.

12. Byrska-Bishop, M., Evani, U.S., Zhao, X., Basile, A.O., Abel, H.J., Regier, A.A., Corvelo, A., Clarke, W.E., Musunuri, R., Nagulapalli, K., et al. (2022). High-coverage whole-genome sequencing of the expanded 1000 Genomes Project cohort including 602 trios. Cell 185, 3426-3440.e19.

13. Hail Team. Hail 0.2. https://github.com/hail-is/hail

14. Laricchia, K.M., Lake, N.J., Watts, N.A., Shand, M., Haessly, A., Gauthier, L., Benjamin, D., Banks, E., Soto, J., Garimella, K., et al. (2022). Mitochondrial DNA variation across 56,434 individuals in gnomAD. Genome Res. 32, 569–582.

15. Dolzhenko, E., Deshpande, V., Schlesinger, F., Krusche, P., Petrovski, R., Chen, S., Emig-Agius, D., Gross, A., Narzisi, G., Bowman, B., et al. (2019). ExpansionHunter: a sequence-graph-based tool to analyze variation in short tandem repeat regions. Bioinformatics 35, 4754–4756.

16. Poplin, R., Chang, P.-C., Alexander, D., Schwartz, S., Colthurst, T., Ku, A., Newburger, D., Dijamco, J., Nguyen, N., Afshar, P.T., et al. (2018). A universal SNP and small-indel variant caller using deep neural networks. Nat. Biotechnol. 36, 983–987.

17. Yun, T., Li, H., Chang, P.-C., Lin, M.F., Carroll, A., and McLean, C.Y. (2021). Accurate, scalable cohort variant calls using DeepVariant and GLnexus. Bioinformatics.

18. Collins, R.L., Brand, H., Karczewski, K.J., Zhao, X., Alföldi, J., Francioli, L.C., Khera, A.V., Lowther, C., Gauthier, L.D., Wang, H., et al. (2020). A structural variation reference for medical and population genetics. Nature 581, 444–451.

19. Chen, X., Schulz-Trieglaff, O., Shaw, R., Barnes, B., Schlesinger, F., Källberg, M., Cox, A.J., Kruglyak, S., and Saunders, C.T. (2016). Manta: rapid detection of structural variants and indels for germline and cancer sequencing applications. Bioinformatics 32, 1220–1222.

20. Rausch, T., Zichner, T., Schlattl, A., Stütz, A.M., Benes, V., and Korbel, J.O. (2012). DELLY: structural variant discovery by integrated paired-end and split-read analysis. Bioinformatics 28, i333–i339.

21. Gardner, E.J., Lam, V.K., Harris, D.N., Chuang, N.T., Scott, E.C., Pittard, W.S., Mills, R.E., 1000 Genomes Project Consortium, and Devine, S.E. (2017). The Mobile Element Locator Tool (MELT): population-scale mobile element discovery and biology. Genome Res. 27, 1916–1929.

22. Klambauer, G., Schwarzbauer, K., Mayr, A., Clevert, D.-A., Mitterecker, A., Bodenhofer, U., and Hochreiter, S. (2012). cn.MOPS: mixture of Poissons for discovering copy number variations in next-generation sequencing data with a low false discovery rate. Nucleic Acids Res. 40, e69.

23. Pedersen, B.S., Layer R., Quinlan, A.R. (2020). smoove: structural-variant calling and genotyping with existing tools. https://github.com/brentp/smoove

24. Dalton, R. (2002). Tribe blasts “exploitation” of blood samples. Nature 420, 111.

25. Correard, S., Arbour, L., and Wasserman, W.W. (2022). Allele Dispersion Score: Quantifying the range of allele frequencies across populations, based on UMAP. BioRXiv.

26. Choudhury, A., Aron, S., Botigué, L.R., Sengupta, D., Botha, G., Bensellak, T., Wells, G., Kumuthini, J., Shriner, D., Fakim, Y.J., et al. (2020). High-depth African genomes inform human migration and health. Nature 586, 741–748.

27. Abouelhoda, M., Faquih, T., El-Kalioby, M., and Alkuraya, F.S. (2016). Revisiting the morbid genome of Mendelian disorders. Genome Biol. 17, 235.

28. Zayed, H. (2016). The Qatar genome project: translation of whole-genome sequencing into clinical practice. Int. J. Clin. Pract. 70, 832–834.

29. Ameur, A., Dahlberg, J., Olason, P., Vezzi, F., Karlsson, R., Martin, M., Viklund, J., Kähäri, A.K., Lundin, P., Che, H., et al. (2017). SweGen: a whole-genome data resource of genetic variability in a cross-section of the Swedish population. Eur. J. Hum. Genet. 25, 1253–1260.

30. Kaja, E., Lejman, A., Sielski, D., Sypniewski, M., Gambin, T., Dawidziuk, M., Suchocki, T., Golik, P., Wojtaszewska, M., Mroczek, M., et al. (2022). The Thousand Polish Genomes-A Database of Polish Variant Allele Frequencies. Int. J. Mol. Sci. 23. https://doi.org/10.3390/ijms23094532.

31. Garcia, M., Juhos, S., Larsson, M., Olason, P.I., Martin, M., Eisfeldt, J., DiLorenzo, S., Sandgren, J., Díaz De Ståhl, T., Ewels, P., et al. (2020). Sarek: A portable workflow for whole-genome sequencing analysis of germline and somatic variants. F1000Res. 9, 63.

32. Tsosie, K.S., Yracheta, J.M., Kolopenuk, J.A., and Geary, J. (2021). We have “gifted” enough: indigenous genomic data sovereignty in precision medicine. Am. J. Bioeth. 21, 72–75.

33. Jagannathan, V., Drögemüller, C., Leeb, T., and Dog Biomedical Variant Database Consortium (DBVDC) (2019). A comprehensive biomedical variant catalogue based on whole genome sequences of 582 dogs and eight wolves. Anim. Genet. 50, 695–704.

34. Zhou, Z.-Y., Li, A., Otecko, N.O., Liu, Y.-H., Irwin, D.M., Wang, L., Adeola, A.C., Zhang, J., Xie, H.-B., and Zhang, Y.-P. (2017). PigVar: a database of pig variations and positive selection signatures. Database (Oxford) 2017.

## REFERENCES (supplementary data)

1. Li, H., and Durbin, R. (2010). Fast and accurate long-read alignment with Burrows-Wheeler transform. Bioinformatics 26, 589–595.

2. Danecek, P., Bonfield, J.K., Liddle, J., Marshall, J., Ohan, V., Pollard, M.O., Whitwham, A., Keane, T., McCarthy, S.A., Davies, R.M., et al. (2021). Twelve years of SAMtools and BCFtools. Gigascience 10.

3. Andrews, S. (2010). FastQC: A Quality Control Tool for High Throughput Sequence Data [Online]. Available online at: http://www.bioinformatics.babraham.ac.uk/projects/fastqc/

4. Pedersen, B.S., and Quinlan, A.R. (2018). Mosdepth: quick coverage calculation for genomes and exomes. Bioinformatics 34, 867–868.

5. “Picard Toolkit.” 2019. Broad Institute, GitHub Repository. https://broadinstitute.github.io/picard/;BroadInstitute

6. Ewels, P., Magnusson, M., Lundin, S., and Käller, M. (2016). MultiQC: summarize analysis results for multiple tools and samples in a single report. Bioinformatics 32, 3047–3048.

7. Poplin, R., Chang, P.-C., Alexander, D., Schwartz, S., Colthurst, T., Ku, A., Newburger, D., Dijamco, J., Nguyen, N., Afshar, P.T., et al. (2018). A universal SNP and small-indel variant caller using deep neural networks. Nat. Biotechnol. 36, 983–987.

8. Yun, T., Li, H., Chang, P.-C., Lin, M.F., Carroll, A., and McLean, C.Y. (2021). Accurate, scalable cohort variant calls using DeepVariant and GLnexus. Bioinformatics.

9. Van der Auwera GA & O’Connor BD. (2020). Genomics in the Cloud: Using Docker, GATK, and WDL in Terra (1st Edition). O’Reilly Media.

10. Hail Team. Hail 0.2. https://github.com/hail-is/hail

11. McLaren, W., Gil, L., Hunt, S.E., Riat, H.S., Ritchie, G.R.S., Thormann, A., Flicek, P., and Cunningham, F. (2016). The ensembl variant effect predictor. Genome Biol. 17, 122.

12. Laricchia, K.M., Lake, N.J., Watts, N.A., Shand, M., Haessly, A., Gauthier, L., Benjamin, D., Banks, E., Soto, J., Garimella, K., et al. (2022). Mitochondrial DNA variation across 56,434 individuals in gnomAD. Genome Res. 32, 569–582.

13. Kirsche, M., Prabhu, G., Sherman, R., Ni, B., Aganezov, S., and Schatz, M.C. (2021). Jasmine: Population-scale structural variant comparison and analysis. BioRXiv.

14. Chen, S., Krusche, P., Dolzhenko, E., Sherman, R.M., Petrovski, R., Schlesinger, F., Kirsche, M., Bentley, D.R., Schatz, M.C., Sedlazeck, F.J., et al. (2019). Paragraph: a graph-based structural variant genotyper for short-read sequence data. Genome Biol. 20, 291.

15. Gardner, E.J., Lam, V.K., Harris, D.N., Chuang, N.T., Scott, E.C., Pittard, W.S., Mills, R.E., 1000 Genomes Project Consortium, and Devine, S.E. (2017). The Mobile Element Locator Tool (MELT): population-scale mobile element discovery and biology. Genome Res. 27, 1916–1929.

16. Robinson, J.T., Thorvaldsdóttir, H., Winckler, W., Guttman, M., Lander, E.S., Getz, G., and Mesirov, J.P. (2011). Integrative genomics viewer. Nat. Biotechnol. 29, 24–26.

